# Nabo – a framework to define leukemia-initiating cells and differentiation in single-cell RNA-sequencing data

**DOI:** 10.1101/2020.09.30.321216

**Authors:** Parashar Dhapola, Mohamed Eldeeb, Amol Ugale, Rasmus Olofzon, Eva Erlandsson, Shamit Soneji, David Bryder, Karlsson Göran

## Abstract

Single-cell transcriptomics facilitates innovative approaches to define and identify cell types within tissues and cell populations. An emerging interest in the cancer field is to assess the heterogeneity of transformed cells, including the identification of tumor-initiating cells based on similarities to their normal counterparts. However, such cell mapping is often confounded by the large effects on total gene expression programs introduced by strong perturbations such as an oncogenic event. Here, we present Nabo, a novel computational method that allows mapping of cells from one population to the most similar cells in a reference population, independently of confounding changes to gene expression programs initiated by perturbation. We validated this method on multiple datasets from different sources and platforms and show that Nabo achieves higher rates of accuracy than conventional classification methods. Nabo is available as an integrated toolkit for preprocessing, cell mapping, differential gene expression identification, and visualization of single-cell RNA-Seq data. For exploratory studies, Nabo includes methods to help evaluate the reliability of cell mapping results. We applied Nabo on droplet-based single-cell RNA-Seq data of healthy and oncogene-induced (MLL-ENL) hematopoietic progenitor cells (GMLPs) differentiating in vitro. Despite a substantial cellular heterogeneity resulting from differentiation of GMLPs and the large transcriptional effects induced by the fusion oncogene, Nabo could pinpoint the specific cell stage where differentiation arrest occurs, which included an immunophenotypic definition of the tumor-initiating population. Thus, Nabo allows for relevant comparison between target and control cells, without being confounded by differences in population heterogeneity.

## INTRODUCTION

It is increasingly accepted that the majority of tumors are organized in cellular hierarchies, where the cancer stem cells or tumor-initiating cells (TICs) drive tumor growth (Clevers, 2011). TICs possess unique stem cell characteristics such as self-renewal capacity, quiescence and drug resistance, which underlie metastasis and relapse. TICs are therefore a critical priority as targets for therapy. The development of improved immune-deficient mouse strains together with refinements to FACS-based protocols for prospective isolation of functionally distinct progenitor populations have paved the way for the definition of the immunophenotype of a plethora of TIC-containing cell populations (Al-Hajj et al., 2003; Boiko et al., 2010; Bonnet and Dick, 1997; Quintana et al., 2008; Schepers et al., 2012; Singh et al., 2004; Wang and Dick, 2005). Even though these efforts have been instrumental in shedding light on TIC biology, including the identification of leukemia-initiating progenitor populations, they have also demonstrated that immunophenotypically defined populations are predominantly heterogeneous. Consequently, many times only a fraction of the cells are relevant for the research aim. Solving the cellular heterogeneity within TIC-containing fractions is particularly critical for molecular characterization and therapeutic target-identification, as transcriptional programs might otherwise be confounded by irrelevant cells.

During the last half a decade, a series of technological breakthroughs have radically improved our ability to quantify transcriptomes at the level of single cells (Hashimshony et al., 2012; Jaitin et al., 2014; Macosko et al., 2015; Picelli et al., 2014; Gierahn et al., 2017; Zheng et al., 2017). Clubbed under the generic term single-cell RNA-Seq (scRNA-Seq), this array of technologies has allowed for detailed analysis of cellular heterogeneity (Zeisel et al., 2015; Tirosh et al., 2016). Indeed, scRNA-Seq has led to the identification of new cell types (Grün et al., 2015; Ramsköld et al., 2012; Villani et al., 2017), deconstruction of developmental programs and lineage hierarchies (Treutlein et al., 2014; Trapnell et al., 2014; Shin et al., 2015; Blakeley et al., 2015; Moignard et al., 2015; Paul et al., 2015), spatial localization of cells in tissues (Achim et al., 2015; Satija et al., 2015), effects of high-throughput gene editing (Dixit et al., 2016) and cell reprogramming (Treutlein et al., 2016). A flurry of analysis methods has followed experimental innovations in this area. Most algorithms and software have focused on certain aspects of scRNA-Seq: data normalization (Vallejos et al., 2017), identification of cell clusters (Andrews and Hemberg, 2018), construction of lineage trajectories (Herring et al., 2018) and identification of differentially expressed genes (Jaakkola et al., 2017). Most recently, data integration from multiple experiments has received necessary attention (Butler et al., 2018; Haghverdi et al., 2018; Kiselev et al., 2018).

Increasing efforts in the genomics area now focus on investigating changes in heterogeneity following a cellular or molecular perturbation. This is of particular interest in cancer research. Here, the effect on cellular heterogeneity by oncogenic transformation or drug treatment has a critical impact for defining TICs, understanding the molecular mechanisms behind therapy resistance, and identifying TIC-specific therapeutic targets. One conceptual strategy to analyze such data is to perform ‘cell mapping’, wherein one of the samples is considered to be a base or reference population (for example a healthy cell population). Individual cells from one or more perturbed populations, called target populations hereon (e.g. tumor cells) are then mapped/projected to the reference population with the objective to identify their most similar counterparts. The current methods of cell mapping (Kiselev et al., 2018), cell alignment (Butler et al., 2018) and batch correction (Haghverdi et al., 2018) rest on the critical assumption that the dissimilarity between the test cells and one or more reference subgroups is smaller than that between at least any one pair of reference subgroups. However, this assumption may not hold true in experimental settings wherein the expression variance of genes responsible for the cellular heterogeneity is smaller than the molecular response to the perturbation. This is often the case when comparing a cancer cell population to its heterogeneous population-of-origin and if cells have been perturbed to alter lineage determining transcriptional networks. Thus, to increase the impact of scRNA-Seq technologies, development of novel bioinformatics tools for improved cell mapping is warranted.

To address these challenges, we here introduce Nabo, a novel computational approach for cross-population cell mapping. Nabo provides a graph-theory based approach to statistically validate mapping and allows integrative comparison of multiple target populations to the same reference population. Using a variety of published as well as in-house generated datasets, we show that Nabo performs equally or better than currently available methods for conventional cell mapping. To demonstrate the power and impact of Nabo for cancer research, we perform scRNA-Seq analysis on the cancer stem cell-containing population from our MLL-ENL mouse model for acute myeloid leukemia (Ugale et al., 2014) and demonstrate how Nabo, unlike current state-of-the art cell-mapping methods, readily identifies leukemia-initiating cells within a heterogeneous population. Thus, Nabo represents a novel tool for relevant analysis of scRNA-Seq data in which a perturbation to an originally heterogeneous population results in a large molecular change to the target cells. As such, Nabo has a critical implementation in cancer research for detection of TICs.

## RESULTS

### Nabo maps cell populations across datasets with high accuracy

The generation of mouse models for leukemia by enforced expression of oncogenic fusion genes has been instrumental for our current conceptual understanding of the origin of cancer stem cells. Interestingly, while the oncogenic targeting of hematopoietic stem cells almost consistently results in leukemic transformation, other progenitor populations also has transformation potential (Eppert et al., 2011; Heuser et al., 2011; Huntly et al., 2004; Krivtsov et al., 2006; Somervaille and Cleary, 2006; Ugale et al., 2014). Thus, the generation of cancer stem cells does not necessarily include high-jacking of the normal stem cells’ molecular machinery, but could also occur due to re-activation of critical stem cell programs in progenitor populations (Eppert et al., 2011; Krivtsov et al., 2006). In fact, we have recently shown that the MLL-ENL fusion gene exclusively transforms heterogeneous progenitor populations with myeloid differentiation potential downstream of the hematopoietic stem cells (Ugale et al., 2014). Here, we used droplet-based scRNA-Seq technology to try to address whether MLL-ENL transformation expands a specific subpopulation within the heterogeneous GMLP population, which would potentially reveal the origin of the leukemic stem cells in MLL-ENL AML.

Purified GMLPs from transgenic mice carrying a doxycycline (dox) inducible MLL-ENL fusion gene were cultured for five days with or without dox and approximately 2,000 cells were subsequently processed for scRNA-Seq using the Chromium droplet-based platform. T-SNE visualization of the scRNA-Seq data (**supplementary figure 1A**) showed that the un-induced (WT) and induced (MLL-ENL) cells divided into two separate clusters, suggesting large molecular differences caused by the expression of the fusion gene. To rule out that this was not simply due to batch effects or other technical artefacts, we used Seurat’s canonical correlation analysis (CCA) based approach of combining datasets. The t-SNE plot of WT and MLL-ENL cells post CCA alignment (**supplementary figure 1B**) did not show any obvious cell clusters and the population structure that was otherwise observed in WT cells alone was lost after CCA alignment. Together these analyses demonstrate the weakness of conventional bioinformatics tools in comparing single-cell RNA-seq data from a heavily perturbed cell fraction to its heterogeneous population of origin. The dramatic molecular effect of the perturbation that a pre-leukemic lesion represents overshadows the gene expression variation that otherwise separates the different subpopulations within the heterogeneous progenitor population, and thus eliminates the advantages of the single-cell dimension from the analysis.

To specifically address these challenges, we designed a novel computational tool for cross-population cell mapping, called Nabo. When using Nabo to perform cell mapping, samples are divided into reference and test populations, where the test cells are projected or mapped over the reference population. Nabo defines a relationship between cells from the reference sample by creating a shared nearest neighbor (SNN) graph (Jarvis and Patrick, 1973). To create an SNN graph, the distance between each pair of cells is calculated based on the expression levels of genes. For each cell, an arbitrary number of cells that have the least distance to the given cell are identified and are regarded as that cells’ neighbors. Subsequently, cells that have common(shared) neighbors are connected to each other and the strength of each connection is determined by the number of shared neighbors (**figure 1A**). After creation of the SNN graph, Nabo maps cells from one or more target samples onto this reference graph by identifying an arbitrary number of the most similar reference cells for each target cell (**figure 1B**). The test cells are then connected to the reference graph by identifying the shared neighbors. Importantly, whether from the same or different samples, the test cells are always mapped independently from each other and hence can never have connections between themselves (see Methods for further details) (**figure 1C**). This feature will circumvent separation between test and reference cells based on large molecular effects due to the specific perturbation. Upon mapping, Nabo provides a ‘mapping score’ to each of the reference cells based on the number and strength of connections that were made to that reference cell from the target cells (**figure 1D**). Thus, the higher the mapping score of a cell, the higher its similarity to the target cell relative to other reference cells. The reference populations can be partitioned into clusters (**figure 1E**) and cluster-wise mapping scores can be identified to ascertain if the test cells were significantly similar to a subgroup of the reference population (**figure 1F**).

**Figure 1:**
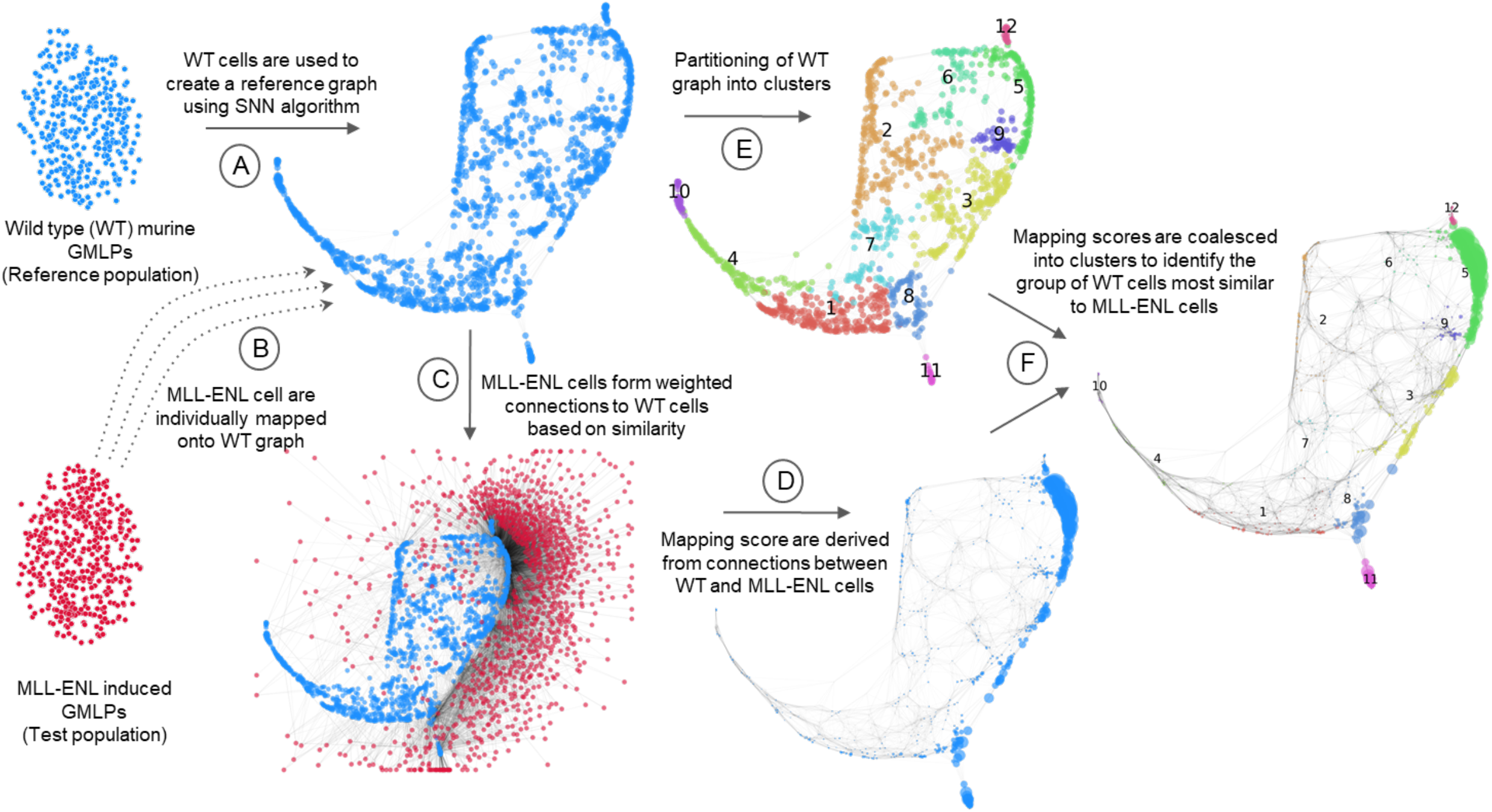
Workflow of cell mapping using Nabo. (**A**) SNN graph of reference population. (**B-C**) Projection of test cells over the reference SNN graph. (**D**) Reference cells sized based on their mapping score. (**E**) Reference cells colored based on their cluster identity. (**F**) Reference cells colored based on cluster identity and sized as per mapping score.

Using scRNA-Seq data from freshly isolated human peripheral blood mononuclear cells (66,000 peripheral blood mononuclear cells) as well as for purified CD19+ B cells, CD14+ monocytes and CD56+ natural killer (NK) cells (Zheng et al., 2017), Nabo correctly mapped cells of known identity onto a more heterogeneous group of cells containing multiple cell types (**figure 2A**). We found that the mapping score of the reference cells was significantly clustered (p < 1e-50; proportions z-test) to the cells expressing the corresponding marker gene (**supplementary figure 2A**). Reference cells expressing CD79A and MEIS1 (B cell marker genes) were mapped by CD19+ B cells with a specificity of 0.976 and 0.995. Similarly, for CD14+ Monocytes: 0.975 (CD14), 0.997 (FTL), 0.988 (LYZ) and for NK cells 0.993 (GNLY) and 0.994 (NKG7) mapping specificity was found. This indicated that Nabo could reliably project cells onto a heterogeneous population with high accuracy. Importantly, Nabo did not require any prior cluster knowledge to perform cell mapping.

**Figure 2:**
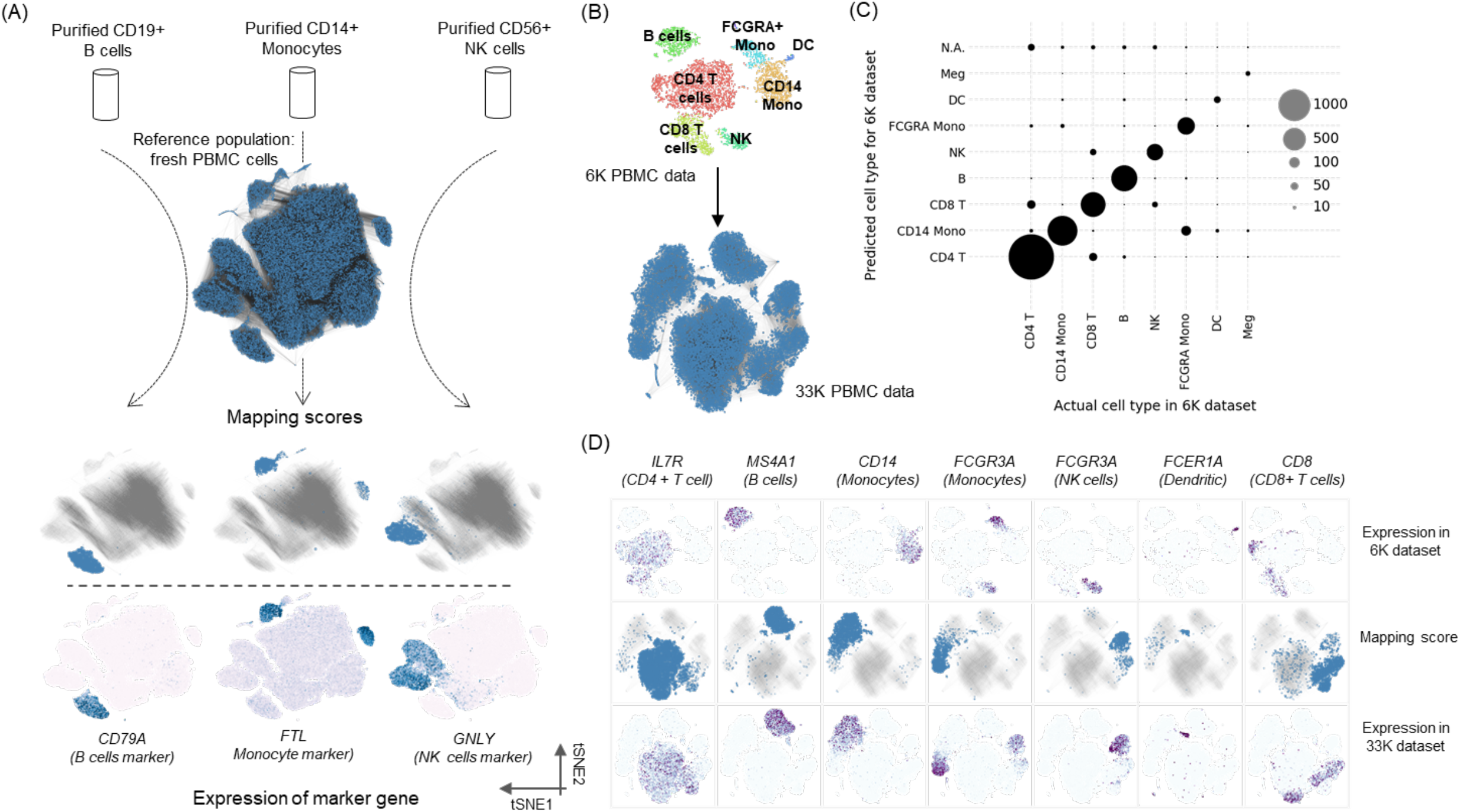
The mapping accuracy of Nabo. (**A**) Mapping of three purified cell populations, CD19+ B cells, CD14+ monocytes and CD56+ NK cells on fresh PBMC cells. In the top t-SNE plot of the reference population, i.e. PBMCs, the cells are connected to each other based on the SNN graph of PBMCs. Middle panel of t-SNE plots shows cells sized based on the mapping scores obtained from mapping of each individual purified population. The cells are colored in blue and the edges in grey. Absence of blue cells indicates that no mapping score was assigned to those reference cells. The bottom panel of t-SNE plots shows the reference cells colored based on expression of marker genes of the respective mapped populations; darker blue color indicates higher expression. (**B**) Mapping of individual clusters of cells from one dataset to another. The t-SNE layout of cells from the 6K dataset has been shown with cells colored based on their cluster identity. The clusters are labeled based on expression of canonical markers. The bottom t-SNE plot shows the reference population, i.e. 33K PBMC dataset. The cells are connected to each other as for the SNN graph. As no clustering was done on this dataset, all cells are colored blue. (**C**) Comparison of cell types in the 6K dataset identified by Nabo and Seurat. Each data point in the scatter plot is sized to indicate number of cells. (**D**) The top panel contains the t-SNE plots of the 6K dataset, with individual cells colored based on the expression of defined marker genes for the indicated cell type. The middle plot depicts a t-SNE layout of the 33K dataset sized based on mapping scores following mapping of the respective 6K cluster. The gray lines indicate the edges in the SNN graph of the 33K dataset. The bottom panel of t-SNE plots shows expression of marker gens in the 33K dataset in same order as the top panel.

To illustrate Nabo’s ability to perform cell identity prediction, we took advantage of two publically available scRNA-seq datasets containing 6,000 (6K) and 33,000 (33K) PBMCs obtained from a healthy donor. We created a reference graph of the 33K dataset, partitioned it into clusters and created a t-SNE layout of the cells for visualization (**supplementary figure 2B**). Using Nabo, we mapped the cells from the 6K dataset cells onto this reference graph, with the objective of identifying the heterogeneity of the 6K cells (**figure 2B**). The cluster identities generated by Nabo correlated to 90.6 % with clusters generated by regular Seurat analysis of the 6K dataset alone (**figure 2C**). Nabo outperformed Random Forest (κ: 0.395), a general classification algorithm, as well as scmap-cell (κ: 0.772) (Kiselev et al., 2018), a single-cell RNA-Seq cell projection tool. Mapping scores were then generated for the 33K cells by individual mapping of each cluster from the 6K data. Importantly, the mappings of distinct clusters from the 6K population were strongly restricted to the 33K cluster, with similar expression levels of marker genes (**figure 2D**). Quantification analysis revealed that 92.34% of all mapping scores were ascribed to cells with correct cluster identity (**supplementary figure 2C**).

Finally, we explored Nabo’s ability to perform cell mapping when reference and target samples are from different studies and use different scRNA-Seq platforms. For this, we used 4 different published scRNA-Seq datasets from pancreatic islets of Langerhans (Baron et al., 2016; Muraro et al., 2016; Segerstolpe et al., 2016; Xin et al., 2016). We chose one of the datasets as the reference population (Baron et al.; InDrop sequencing) and, using Nabo, constructed an SNN graph of the same dataset and calculated a force directed layout of the graph for visualization (**figure 3A**). After mapping the other three datasets onto this reference graph (**figure 3B**), we found that Nabo was able to correctly map 69.3% (Xin et al, C1-IFC SMARTer platform (κ: 0.687)), 63.79% (Muraro et al., CEl-Seq2 platform (κ: 0.733)) and 88.36% (Segerstolpe et al SMART-Seq2 platform (κ: 0.927)) of the target cells to their appropriate cluster type as defined by the reference dataset (**figure 3C**). In comparison, scmap-cell correctly mapped 78.75% (κ: 0.665), 81.32% (κ: 0.764) and 50.65% (κ: 0.45) of cells in the three datasets. Thus, as per the Cohen’s kappa scores, Nabo performed equally well- or better than scmap-cell in cell mapping using data obtained from different platforms. Together, these experiments demonstrate that Nabo performs equally or better than currently available methods for several different aspects of conventional cell mapping.

**Figure 3:**
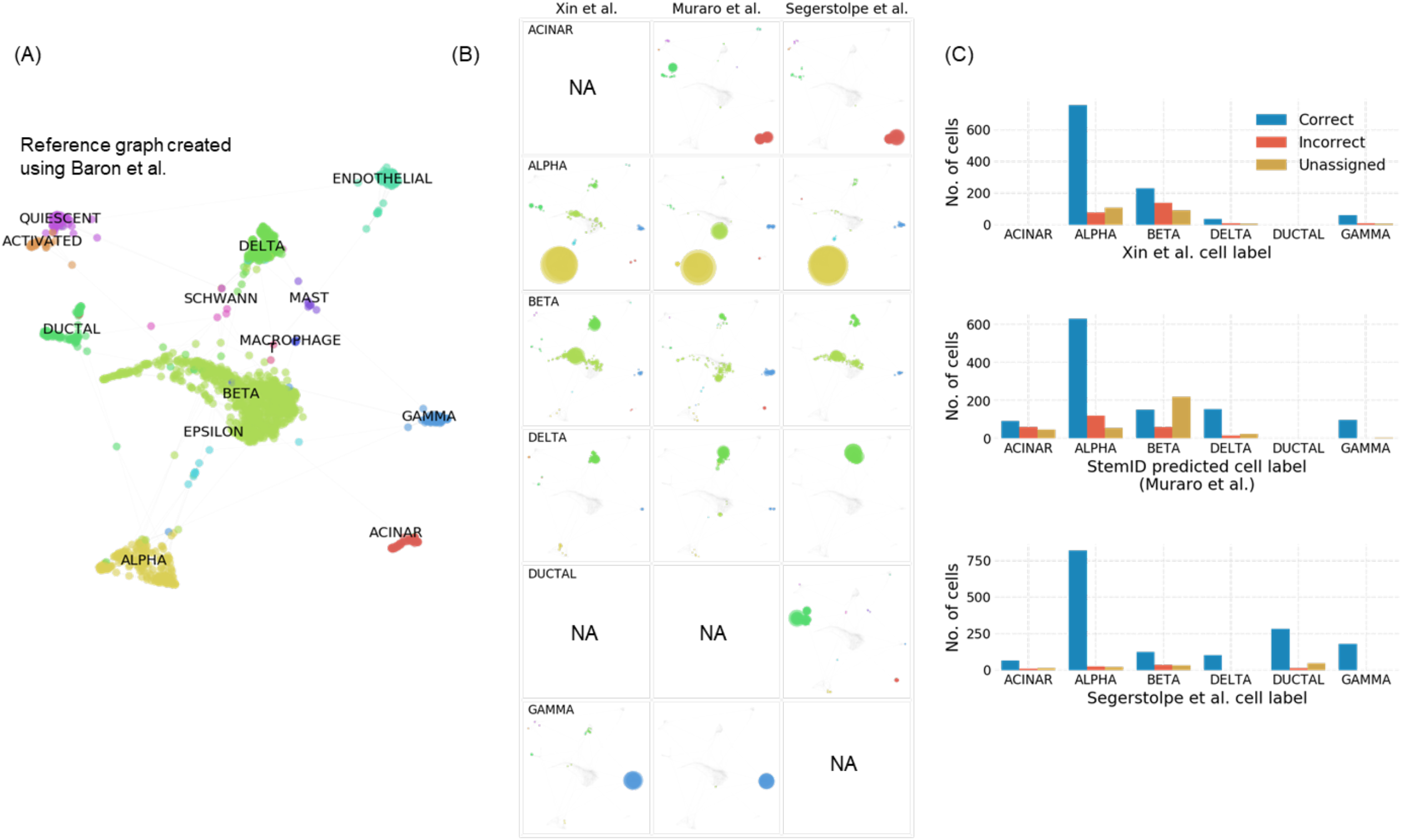
Mapping of pancreatic cells across datasets. (**A**) A force directed layout of pancreatic cells obtained from Baron et al. showing clusters of cells labelled based on marker expression. This cell population was used as reference population for mapping from other datasets. (**B**) Each panel shows a force directed layout of reference cells, with cells scaled in size based on mapping scores obtained by mapping of specific cell types from a given study. NA panel indicate that the cell type was not present in that study. (**C**) Barplots showing the number of cells whose cell type was predicted either correctly, incorrectly or remained unassigned with respect to cell types reported in the original study.

### Identification of MLL-ENL leukemia-initiating cells using Nabo

Having an improved mapping tool at hand, we used Nabo on our scRNA-Seq data to compare the heterogeneity of MLL-ENL-transformed and WT GMLPs. A shared nearest neighbor graph of WT cells was created and partitioned into 12 clusters (**figure 4A**). MLL-ENL cells were projected onto this WT graph, providing a mapping score for each WT cell that signifies the reference cell’s similarity to the MLL-ENL cells (**figure 4B**). MLL-ENL cells were classified as members of either one of the WT graph clusters or they remained unassigned if enough evidence to perform assignment was lacking. Intriguingly, WT cells with high mapping score were significantly (p<1e-6; Chi-squared test) closely connected on the graph where cluster 5 of WT cells had the absolute highest average mapping score (5.02) followed by cluster 11 (1.89) (**figure 4C**). Of all MLL-ENL cells that were assigned to a cluster, 70.44% were assigned to cluster 5. In contrast, scmap-cell was unable to assign the vast majority (98.73%) of MLL-ENL cells to any cluster, while scmap-cluster assigned most of the MLL-ENL cells (77.5%) to cluster 6. However, with increased stringency (see Methods), scmap-cluster failed to assign 81.56% of MLL-ENL cells to any cluster (**figure 4D**). To test how the results of each tool were dependent on the clustering itself, we used Seurat to perform clustering of the reference population (**supplementary figure 3A-B**) and then predicted the identity of MLL-ENL cells again. Even with a different partitioning of the cells, Nabo, under two different stringency settings, assigned 56.18% and 45.6% of MLL-ENL cells to Seurat’s cluster 5, which mostly comprised of the same cells as the cluster 5 from the SNN graph (**figure 4A**). Using this strategy, both scmap-cell and scmap-cluster (with high stringency settings) left 69.29% and 69.96% cells unassigned. Scmap-cell however, did assign 20.98% of cells to Seurat’s cluster 5. With default settings, scmap-cluster assigned 45.51% of cells to cluster 5. Interestingly, scmap-cluster assigned only 1 cell to cluster 10 and no cell to cluster 11, which combined constituted the cells from the SNN cluster 6 (**figure 4A**), to which scmap-cluster had assigned 916 cells (77.5%) (**supplementary figure 3C**). The mapping performed by Nabo clearly indicates that the MLL-ENL cells bear a strong relative preference for cells that are from cluster 5. We also found that Nabo, unlike scmap, consistently predicted the association of MLL-ENL cells to WT clusters across different cluster partitioning.

**Figure 4:**
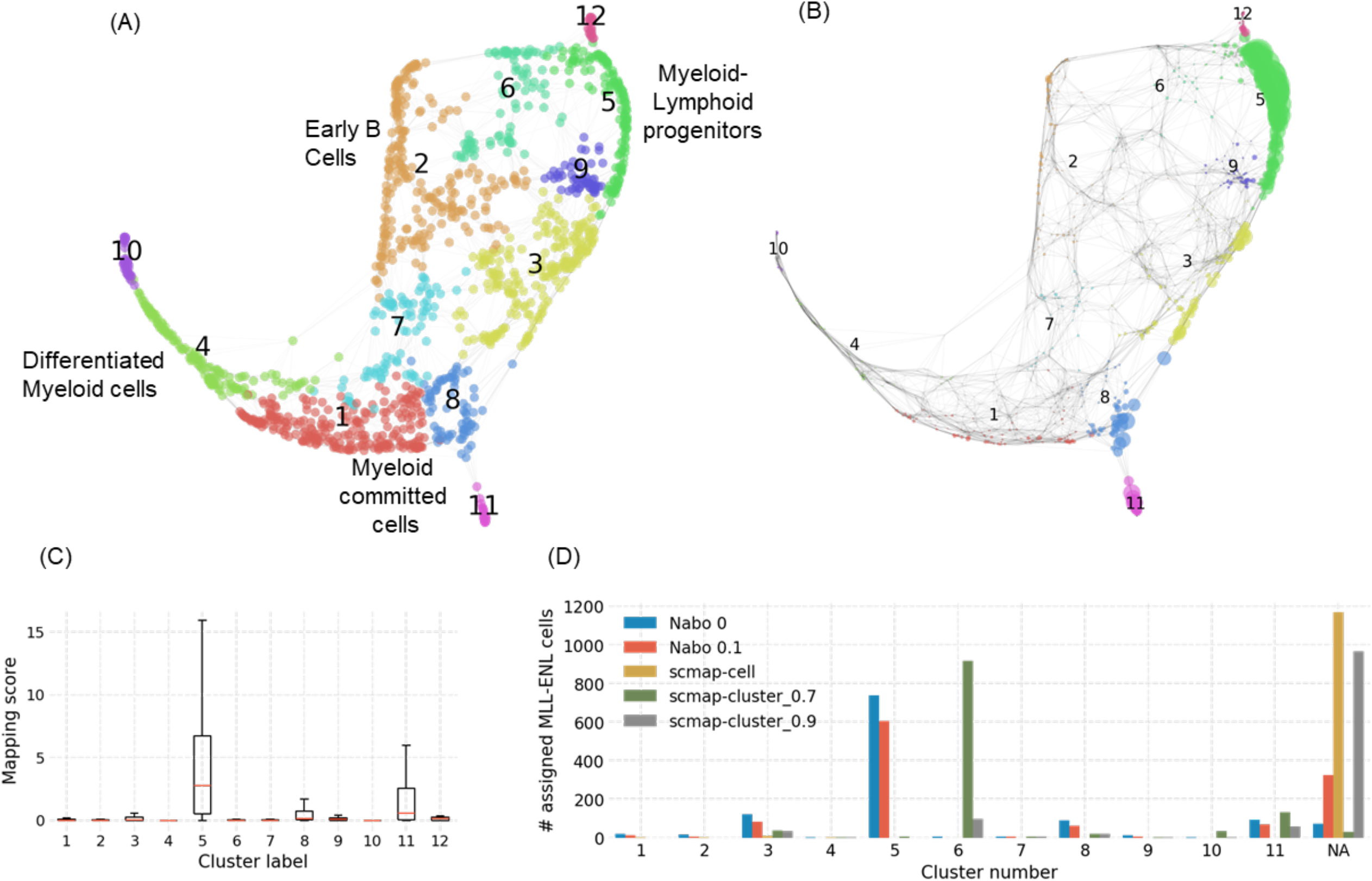
Mapping of MLL-ENL expressing GMLPs. (**A**) A force directed layout of an SNN graph depicting normal GMLPs, with clusters assigned to numerals and colors. The different cell stages inferred from gene expression patterns is indicated. (**B**) SNN graph of normal GMLPs where cells have been sized based on their mapping score obtained after projection of MLL-ENL induced GMLPs. (**C**) Boxplots showing the distribution of mapping scores across the clusters. The red line in each box indicates the median value. (**D**) Barplots showing the number of MLL-ENL cells that were assigned to each reference graph cluster. The minimum weight for assignment using Nabo was set at either 0 or 0.1 (default). In case of scmap-cluster, the weight fraction was set either at 0.7 (default) or 0.9.

As a control to address if the results obtained from Nabo-generated mapping were due to technical biases such as -cell-cycle effect or sequencing depth, we used low variance genes (LVGs) rather than high variance genes to create the reference graph and perform the mapping. We visualized the mapping scores on LVG reference graph using the same layout as for the actual reference graph. This revealed a substantially more even distribution of cells across the graph (**supplementary figure 4A**). Additionally, the distribution of LVG mapping scores across the clusters was relatively uniform, with cluster 5 having an insignificantly higher mapping score distribution than others (**supplementary figure 4B**). However, in absolute terms, the mean mapping score in cluster 5 was 33.78 times lower in LVG mapping compared to original (HVG) mapping. Also, using LVG mapping, Nabo failed to assign 92.89% of cells to any cluster (**supplementary figure 4C**). To investigate the robustness of the mapping, we prevented MLL-ENL cells from mapping to reference cells that had received a mapping score higher than 1 in the actual mapping (10.75% cells). If WT cells with high mapping scores were not significantly more similar to MLL-ENL cells than to the rest of the WT cells, then in this control setup, called blocked mapping, a subset of these cells will receive similarly high mapping scores. While 58 of the reference cells received a mapping score higher than 5, only one cell passed this threshold in blocked mapping (**supplementary figure 4D**). Under these blocked mapping conditions, cluster 9 received the highest mapping scores (**supplementary figure 4E**). However, this was still 5.86 times lower than the mean mapping score of cluster 5 from the actual mapping (figure 4c). Also, 64.13% of cells were not assigned to any cluster by Nabo’s classifier (**supplementary figure 4F**), which was 2.33 times higher than observed in the actual mapping (**figure 4C**). Overall, these two controls indicated that the mapping of MLL-ENL on WT cells was significant and non-trivial. As these controls are critical and useful methods for evaluating mapping results in exploratory studies, they have been assigned to Nabo for easy inclusion in an analysis workflow. Within Nabo, we have also included a heuristic to identify mapping specificity of each target node. Mapping specificity of a node will be higher if all the reference nodes it connects to are also close to each other. When applied to MLL-ENL cells, we found that the cells that projected to cluster 5 and cluster 11 (**figure 4A and 4C**) had higher average mapping specificity than other MLL-ENL cells (**supplementary figure 5A**).

These mapping specificities can also be viewed from a reference cell’s perspective, by averaging the mapping specificity of all the target cells that map to a given reference cell (**supplementary figure 5B**).

To define the identity of WT cells in cluster 5, to which an overwhelming majority of the MLL-ENL cells had mapped, we used Nabo to identify marker genes specifically expressed within this cluster. The cumulative expression of these marker genes were then cross-referenced to known cell types from bulk transcriptome datasets obtained from the BloodSpot database (Bagger et al., 2016). Using the annotations of each cluster, a myeloid and a lymphoid differentiation trajectory could be inferred in the WT graph (**figure 4A**). Interestingly, within this trajectory, the MLL-ENL-mapped cluster 5 as well as cluster 9 represented subpopulations with the most primitive molecular signatures identified as LMPP- and Pre-GM-like, respectively (**supplementary figure 6**). This observation is in concordance with previous reports that suggest that overexpression of oncogenes in pluripotent/multipotent cells can prevent primitive cells from terminally differentiating and subsequently develop into tumors (Cozzio et al., 2003).

To further investigate the gene signature of ‘MLL-ENL like WT cells’, we focused only on a subset of cells within cluster 5 that received a mapping score greater than 1 (92 cells) and compared them to their nearest 89 cell neighbors (see Methods) (**supplementary figure 7A**). We found 96 genes with significantly higher expression (adjusted p-value < 0.05) in MLL-ENL like WT cells, including FLT3, BCL2, SOX4 and AFF3 (**supplementary figure 7B**). Many of these genes were not just higher in test cells compared to control cells, but also when compared to rest of the cells on the graph (**supplementary figure 7C**). Interestingly, WT cells with high mapping scores demonstrated a primitive gene expression signature when compared to the rest of WT cells (**supplementary figure 8A**), but a more differentiated signature compared to MLL-ENL cells that mapped to these WT cells (**supplementary figure 8B**). On the other hand, MLL-ENL cells displayed a higher expression of genes that are normally expressed in hematopoietic stem cells (**supplementary figure 8C**). These results strongly indicate that MLL-ENL induction perturbs a primitive subpopulation of GMLPs and induces a gene expression program that is normally found upstream in the lineage hierarchy.

To further verify that the MLL-ENL induced cells were actually in a less differentiated state than the WT GMLPs, we chose to map the MLL-ENL-transformed GMLPs against total cKit+ BM cells (Säwen et al., 2018), which consists of a wider range of hematopoietic progenitors (**supplementary figure 9A**). First, we mapped 500 highly purified hematopoietic stem cells (HSCs) (LSK CD150+ CD48-; (Säwen et al., 2018)) to the reference graph (**figure 5A**). Interestingly, these 500 cells mapped almost exclusively to a rare group of 7 cells within the cKit+ reference graph, demonstrating the stringency of Nabo and establishing the location of the most primitive cell type within the trajectory of the graph. We then mapped the WT GMLPs and MLL-ENL induced GMLPs on the cKit+ cells (**figure 5B-C**). To quantify the differentiation state of the reference cells, we assigned the differentiation potential for each cell (Weinreb et al., 2018) (**figure 5D**) and inferred a pseudo-time axis of the data (**figure 5E**). This revealed that the sorted HSCs were most upstream, followed by MLL-ENL induced GMLPs and then WT GMLPs (**figure 5E**), thereby confirming our earlier observations (**figure 5A-C**). When annotating the cell clusters of the cKit+ reference population, we found that WT GMLPs mapped cells had signatures of GMPs and differentiated granulocytes, while MLL-ENL mapped cells had highest similarity to gene signatures of preGMs and LMPPs (**supplementary figure 9B**). Thus, Nabo is a powerful tool to predict the identity of TICs from scRNA-Seq data.

**Figure 5:**
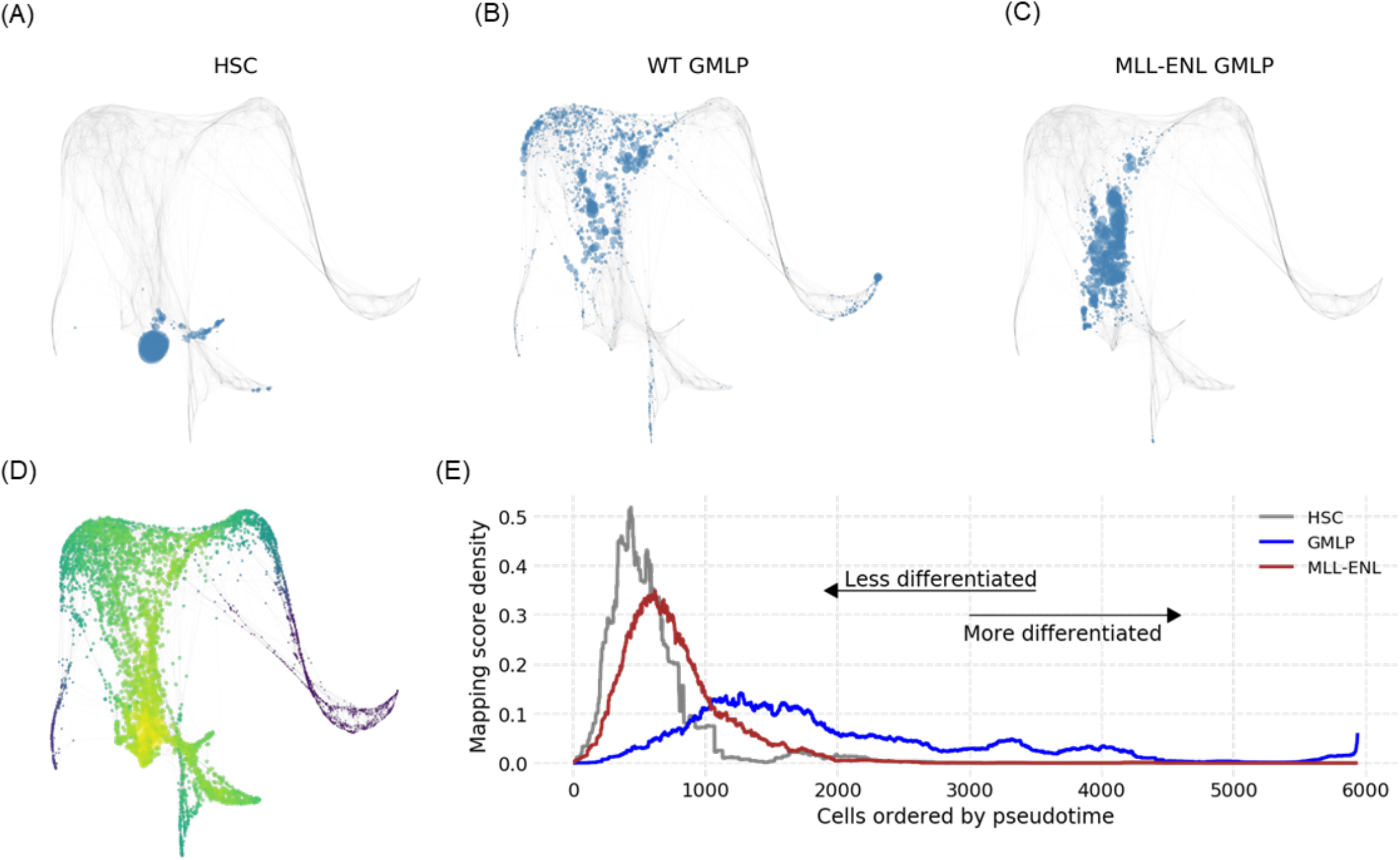
Assessing differentiation of MLL-ENL cells. Mapping of (**A**) purified HSC (**B**) WT GMLPs (**C**) and MLL-ENL induced GMLPs on the SNN graph of cKit+ cells. Cells have been size scaled proportional to the mapping score of respective target population. (**D**) The inferred differentiation potential c-kit+ cells. (**E**) The differentiation potential of cells is shown on the x axis and the y axis shows the average mapping score on cells grouped into bin sizes of 100 cells.

### Nabo has a broad implementation for mapping perturbed populations of different cell systems

Similar to oncogenic transformation, overexpression or deletion of transcription factors may cause large-scale transcriptional changes and disrupt the heterogeneity of a cell population. Such effects will create substantial differences between the normal and perturbed cells, resulting in their independent clustering. To evaluate if Nabo would be a useful tool for these kinds of experimental settings, we used a scRNA-seq dataset comparing differentiating WT murine embryonic stem cells (ESCs) with ESCs deficient for the transcription factor YY1 (Weintraub et al., 2017).

Using tSNE visualization, it was previously observed that YY1-cells clustered away from the YY1+ cells, resulting in a loss of single-cell resolution (Weintraub et al., 2017). In an attempt to compare the heterogeneity between YY1+ and YY1-cells, we used Nabo and created a reference graph for YY1+ cells that was visualized using a force directed layout after partitioning the graph into six clusters (**figure 6A**). After mapping the YY1-cells (**figure 6B**), we found that the mapping scores were mainly concentrated in the clusters corresponding to clusters 4, 5 and 6 (**figure 6C**). Visualizing marker genes for pluripotency and primary germ layers **(figure 6D**) revealed that clusters 5 and 6 were dominated by endodermal and mesodermal molecular signatures, respectively. As the mapping scores were largely low or absent in cells with pluripotency and ectodermal lineage marker expression, we inferred that YY1 depletion in murine ESCs either results in accelerated differentiation towards mesodermal and endodermal lineages, or in the loss of pluripotent and ectodermal progenitor cells. This finding is consistent with the notion that YY1 is critically important for ectodermal development (Satijn et al., 2001) and demonstrate the utility of Nabo to define cellular heterogeneity following different types of perturbation in a broad range of cell types.

**Figure 6:**
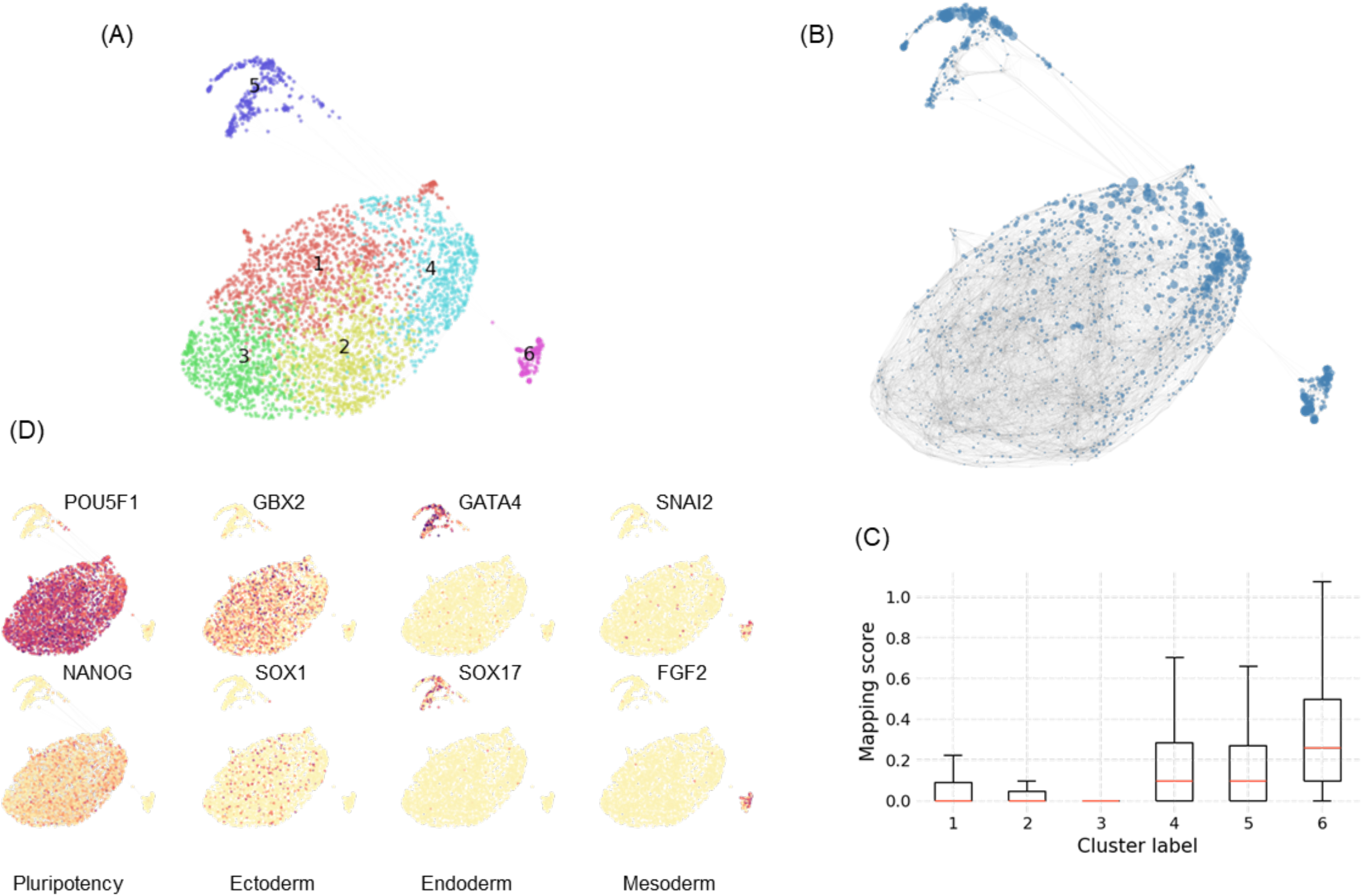
Nabo interrogation of the effects of YY1 depletion on murine embryonic stem cells. (**A**) SNN graph of murine embryonic stem cells (mESCs) where cells have been colored based on their cluster identity. (**B**) SNN graph of mESCs, where node size has been scaled proportional to the mapping scores obtained on projection of YY1 depleted mESCs. (**C**) Distribution of mapping scores across the mESC clusters. (**D**) SNN graph of YY1+ ES cells with expression of marker genes for each germ layer as well as pluripotency associated genes highlighted.

## DISCUSSION

Recent advances in RNA sequencing methodology has allowed for measurements of global gene expression programs in individual cells as a proxy for their function. Thus, scRNA-Seq experiments offer an unprecedented possibility to dissect cellular heterogeneity in tissues or purified cell-fractions, that is now extensively used to visualize cellular hierarchies and compositions throughout the entire human body. Most approaches for analysis of scRNA-seq data have focused on statistical methods to discriminate cells by clustering into groups or trajectories. However, for scRNA-Seq to become useful when approaching changes in heterogeneity during situations of major perturbations of molecular programs, computational methods that allow for comparison of relevant information between datasets are critical. One way to integrate such data is through cell mapping, which allows identification of cell-cell relationships even when the data is spread across experiments, alternative sequencing platforms and studies. With Nabo, we aimed to develop an accessible and interpretable platform to perform cell mapping. The control datasets indicated that Nabo has equal or improved accuracy than existing methods in conventional classification of target cells based on reference cell clusters. Nabo supplements two other recently published algorithms that aim to integrate scRNA-Seq datasets, mmCorrect (mutual nearest neighbors correction) (Haghverdi et al., 2018) and Seurat’s CCA (canonical covariate analysis) based cell alignment (Butler et al., 2018). Both of these algorithms are geared towards removal of batch effects between the datasets. The implicit assumption made by these algorithms is that the differences between the populations being compared is mostly technical in nature. Nabo is not based on such assumptions. Instead, Nabo allow for relevant comparison of heterogeneity between populations even if they have large differences in gene expression profiles due to a biological component, such as overexpression of an oncogene. MNNcorrect is more useful when a user wants to obtain batch corrected expression values, while Seurat’s CCA can be useful in scenarios where the user wants an integrated low-dimensional embedding of two or more datasets. Both of these approaches, however, lead to changes of cell embeddings of the reference population for each new population/sample included. For example, the t-SNE layout of cells will change when new samples are included. Nabo solves this by providing a quantitative measure that can be ascribed to each cell of the reference sample. In this way, no matter which and how many target populations are projected, the reference cells can always be visualized in their original space.

In cell mapping and other predictive analytics approaches that use scRNA-seq data, one major challenge has been the validation of the results. We demonstrated two innovative approaches that Nabo uses to evaluate the mapping reliability that we believe could be particularly useful for experiments where little prior knowledge about the mapping exists. The implementation of Nabo was done with two objectives: high accuracy and low memory requirements, so that the software can be run on ordinary desktops and laptops with modest hardware requirements. The tradeoff for this implementation was a runtime that increases quadratically with cell number. We were able to process PBMC data with 66,000 cells within 6hrs with just 8 GB of RAM required. Most scRNA-Seq experiments carried out today are well within that range.

Importantly, we also demonstrated an exclusive capacity of Nabo to ascribe mapping scores to reference cells. This allows for quantitative assessment of mapping and subsequent identification of specific groups of cells in the reference population that are similar to the test cells, which is advantageous when trying to determine the exact identity of TICs. Unlike current methods, Nabo is designed to ignore the massive transcriptional changes associated with a strong perturbant such as the onset of a strong oncogene and allow for cell mapping entirely based on expression of molecular signatures associated with heterogeneity. Using Nabo on scRNA-Seq data acquired from the TIC-containing GMLP population from our MLL-ENL mouse model of AML, we could identify a distinct target population characterized by a primitive and multipotent molecular signature. Interestingly, this population could be discriminated from other more differentiated GMLP populations by the expression of fms-like tyrosine kinase 3 (Flt3). Together, these results validate the usefulness of Nabo as a tool for scRNA-Seq analysis of tumor populations with the aim of defining the changes in heterogeneity caused by oncogenic transformation and subsequent TIC identification.

## METHODS

### Nabo overview

Nabo uses HDF5 file format to store the data on the disk. The data is stored in gene-wise and cell-wise manner to allow quick subselection across both axes. For mapping, the reference dataset is first normalized (library size normalization) and the selected features are subjected to standard scaling. The data is normalized and scaled on the fly as it is being loaded from the disk. The scaled data is subjected to PCA reduction using an out-of-core (incremental) implementation of PCA. The Euclidean distances are computed between each pair of cells in the PCA space to identify k-nearest neighbor of each reference cell. The shared nearest neighbors are identified for each pair of KNN neighbors and if they have non-zero shared neighbors then an edge is added between the two cells with weight equal to the ratio of number of shared neighbors (*s*) to s – maximum possible shared neighbors. The cells to be projected are too library scale normalized but their features are scaled as per the mean and standard deviation of features in the reference dataset. This scaled target data is then projected into the PCA space trained on reference data. The distance of each target cell to every reference cell is calculated using a modified Canberra metric. The metric is modified such that if in a given dimension the distance between target (*t*) and reference (*r*) cell value is greater than *f* * *r*, where f is a predefined factor between a range of 0-1, then distance is set as 1 (highest value). This means that if the target cell has a very high value in a given dimension, then that dimension would automatically cause the distance to saturate.

### Mapping score calculation

Mapping score is calculated using the ‘*get_mapping_score*’ function of the *Graph* class. The score for a reference cell is calculated by calculating the weighted sum of all of its incoming projections. Mapping scores are always normalized to the number of projected cells, to allow comparison between two mapping score distributions as long as the other parameters remain similar.

### Classification of cells

The classification of target cells is performed by the ‘*classify_targets*’ function of the *Graph* class. The target cells are classified to one of the reference clusters using voting method. The projections (edges) of a given target cell to reference cells are grouped based on the cluster identity of the reference cells. A weighted sum of projections is calculated for each of the groups. If a single group has a weighted sum that is higher than predefined threshold (default: 50% of total weighted sum), then the cell is classified to that cluster otherwise the target cell remains unassigned to any cluster. As an additional filter, projections can be discarded during calculation of cluster-wise weighted sums based on individual weight of each edge connection between a target cell and a reference cell. Furthermore, a target cell can directly be classified as unassigned if it has fewer projections than a pre-set cutoff. These two tunable parameters ensure that users have fine control on classification accuracy.

### Differential gene expression calculation

Mann-Whitney U test (as available in the scipy.stats package in Python) is used to identify genes that are differentially expressed between two groups of cells. P values are corrected for multiple hypotheses testing using the Benjamini/Hochberg method (as implemented in the statsmodels package). If the number of cells in the control are higher than the number of cells in the test group, the same number of cells as in the test group are selected from the control group. Such subselection of cells is performed after sorting the control cells in order to retrieve top *nc* values (from highest to lowest), where *nc* in number of test cells. This strategy helps improve the specificity of the results. To identify the genes driving a mapping specificity, reference cells with mapping score higher than a given threshold are selected (test cells); thereafter all the cells that are at the node distance of *n* (defined by the user) from the test cells, in the reference graph, are marked as control nodes against which genes upregulated in test nodes are identified. The higher values of *n* will cause identification of larger differences in transcriptomes of mapped and unmapped cells, while smaller values of *n* will highlight the smaller differences between the control and test cells.

### Hematopoietic gene signature identification

Gene sets were queried against the ‘normal mouse hematopoiesis’ dataset that contains cell types from the BloodSpot database (Bagger et al., 2016). The median value of the gene set in each cell type was determined and these values were min-max scaled across cell types. The results were visualized as area plots in polar coordinates to enable quick assessment for presence of cell type signature in a gene set. +/−1 standard deviation is also calculated by usage of biological replicate data of cell types and also visualized to provide a quick assessment of noise in the results.

### Data processing

All the datasets were subjected to cell filtering, HVG identification, Louvain clustering, UMAP and/or tSNE embedding generation and marker gene identification; all using Seurat version 3.0.1. Parameters used of each individual dataset can be found in the online repository. The HVGs identified by Seurat were used to perform reference graph generation of the respective datasets. The same list of HVGs was also used to train classifiers and scmap.

### Single-cell RNA-Seq of in vitro cultured GMLPs and MLL-ENL induction

Granulocyte-macrophage-lymphoid progenitors (GMLPs) were enriched from BM of WT and iMLL/ENL mice by depletion of mature cells using biotinylated antibodies against lineage markers (CD4, CD8a, B220, CD11b, Gr1and Ter-119) and anti-biotin conjugated magnetic beads, according to manufacturer□s instructions (Miltenyi Biotech, Germany). GMLPs were sorted as Lin-Sca1+c-Kit+CD48+CD150- on a FACS Aria II or III cell sorter (Becton Dickinson, San José, CA). Propidium iodide (Invitrogen, Carlsbad, CA) was used to exclude dead cells. The 10,000 sorted GMLPs were maintained in OptiMEM (Invitrogen, Carlsbad, CA) supplemented with 10% FCS, 0.1 mM β-mercaptoethanol (Invitrogen, Carlsbad, CA), 1x Penicillin/Streptomycin (Invitrogen, Carlsbad, CA), SCF (10 ng/ml), IL3 (5 ng/ml), G-CSF (5 ng/ml) (all from Peprotech Inc., Rocky Hill, NJ) and 1 μg/ml doxycycline (Sigma-Aldrich, St. Louis, MO). After 4 days, cells were harvested and subjected to single cell (SC) RNA sequencing. ScRNA-Seq data was generated on the 10X platform (10X Genomics) according to the manufacturer’s instructions.

Antibody clones and suppliers are as follow:

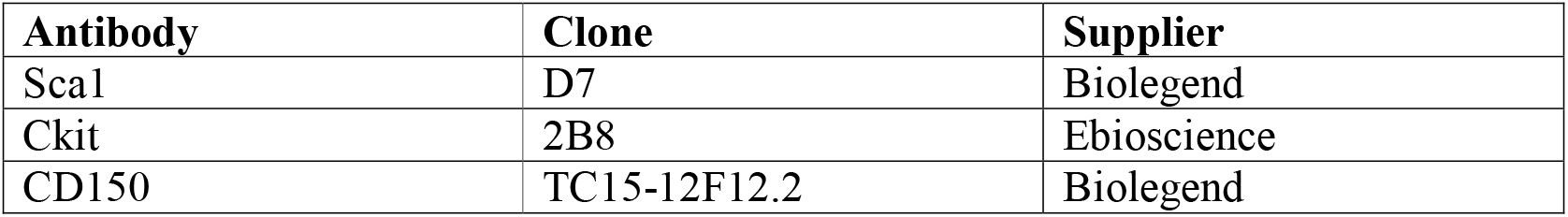
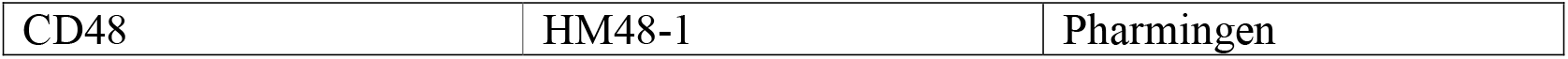

## Data and code availability

All code used to analyze the data and generate the figures can be found here: http://github.com/parashardhapola/nabo_manuscript. This repository also contains the count matrices of the datasets in MTX or CSV format. Nabo format HDF5 files can also be found in the same repository. Source code of Nabo is available here: http://github.com/parashardhapola/nabo. API and tutorials for usage of Nabo can be found here: nabo.readthedocs.io. YY1 dataset was downloaded from GSE103574 (samples: GSM2774584 and GSM2774585). Pancreatic cells datasets were downloaded from these GEO repositories: GSE84133 (Baron et. al.), GSE85241 (Muraro et. al.), GSE81608 (Xin et .al.). Data for Segerstolpe et. al. was downloaded from Array Express archive E-MTAB-5061. PBMC 68K, 33K and 6K cell dataset was obtained from 10x genomics data portal as count matrices (MTX format) generated using Cell Ranger version 1.1.0. Data for murine HSCs and cKIT+ cells was obtained from GSE122473.

---

**Supplementary figure 1:**
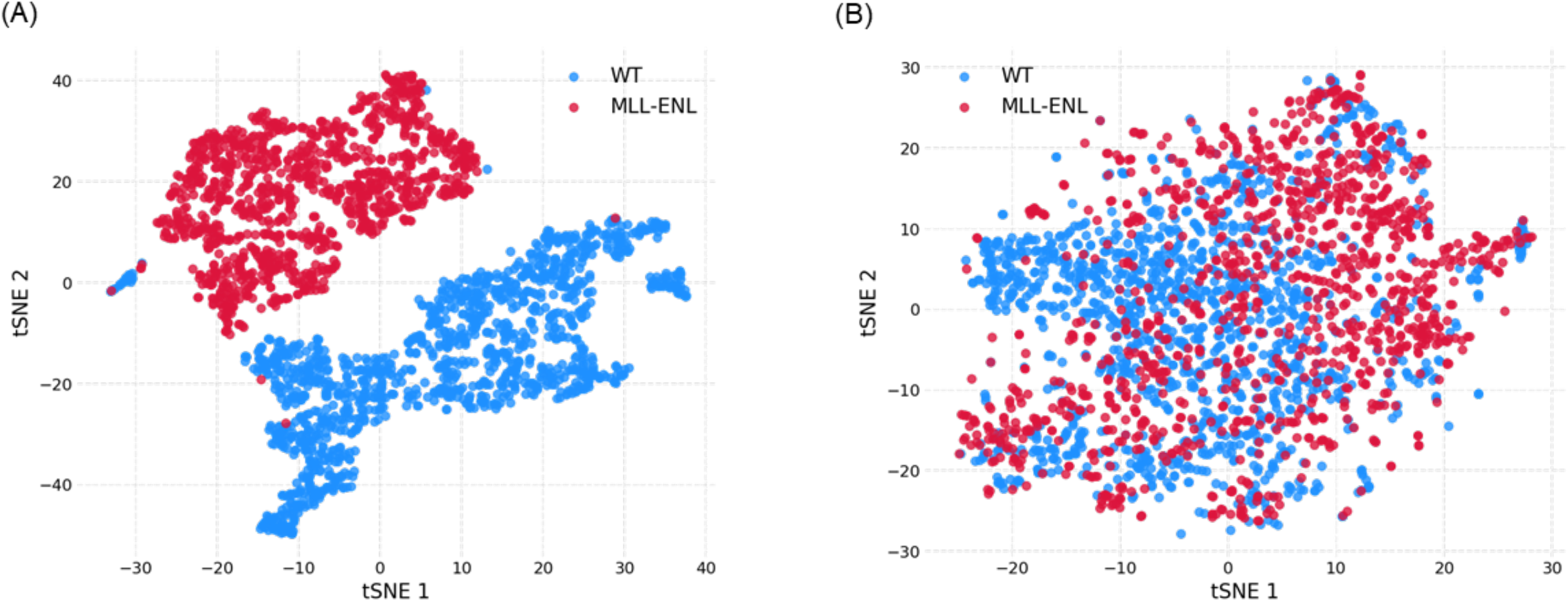
(**A**) t-SNE plot showing cells from WT GMLPs (in blue) and MLL-ENL induced GMLPs (in red) as separate clusters. (**B**) t-SNE plot of WT and MLL-ENL induced GMLPs obtained after applying Seurat’s CCA algorithm.

**Supplementary figure 2:**
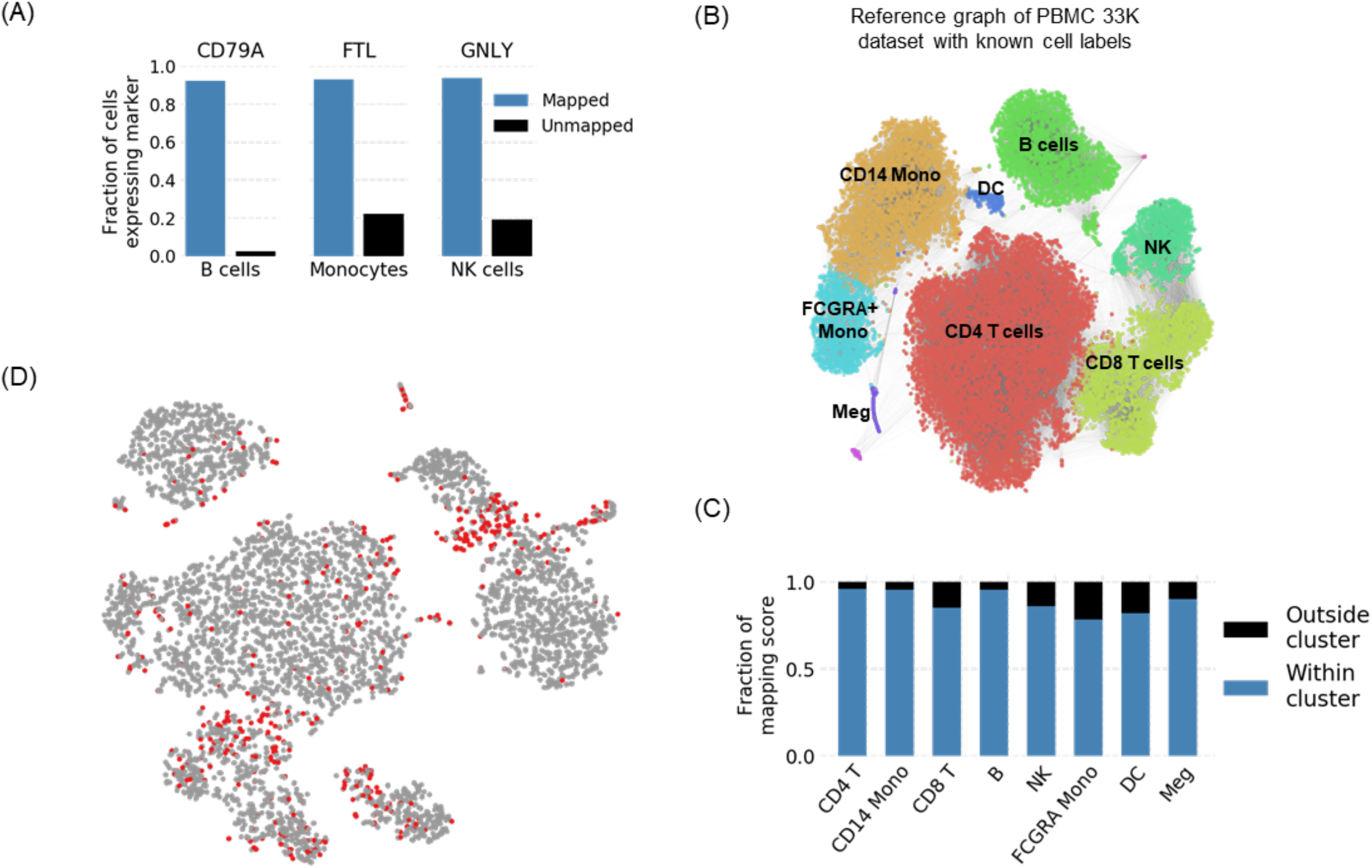
(**A**) The fraction of either mapped or unmapped cells expressing the given marker gene. The mapping was performed using the indicated cell population in the PBMC dataset. (**B**) Reference graph of the 33K data set, with cell labels obtained post clustering using Seurat. Cell labels were placed based on the expression of canonical marker genes. (**C**) Fraction of mapping scores that was either within or outside the 33K dataset Seurat clusters upon mapping of each corresponding 6K cluster. (**D**) A t-SNE visualization of the 6K PBMC dataset showing the cells (in red) whose cluster identity was differently predicted by Nabo and Seurat.

**Supplementary figure 3:**
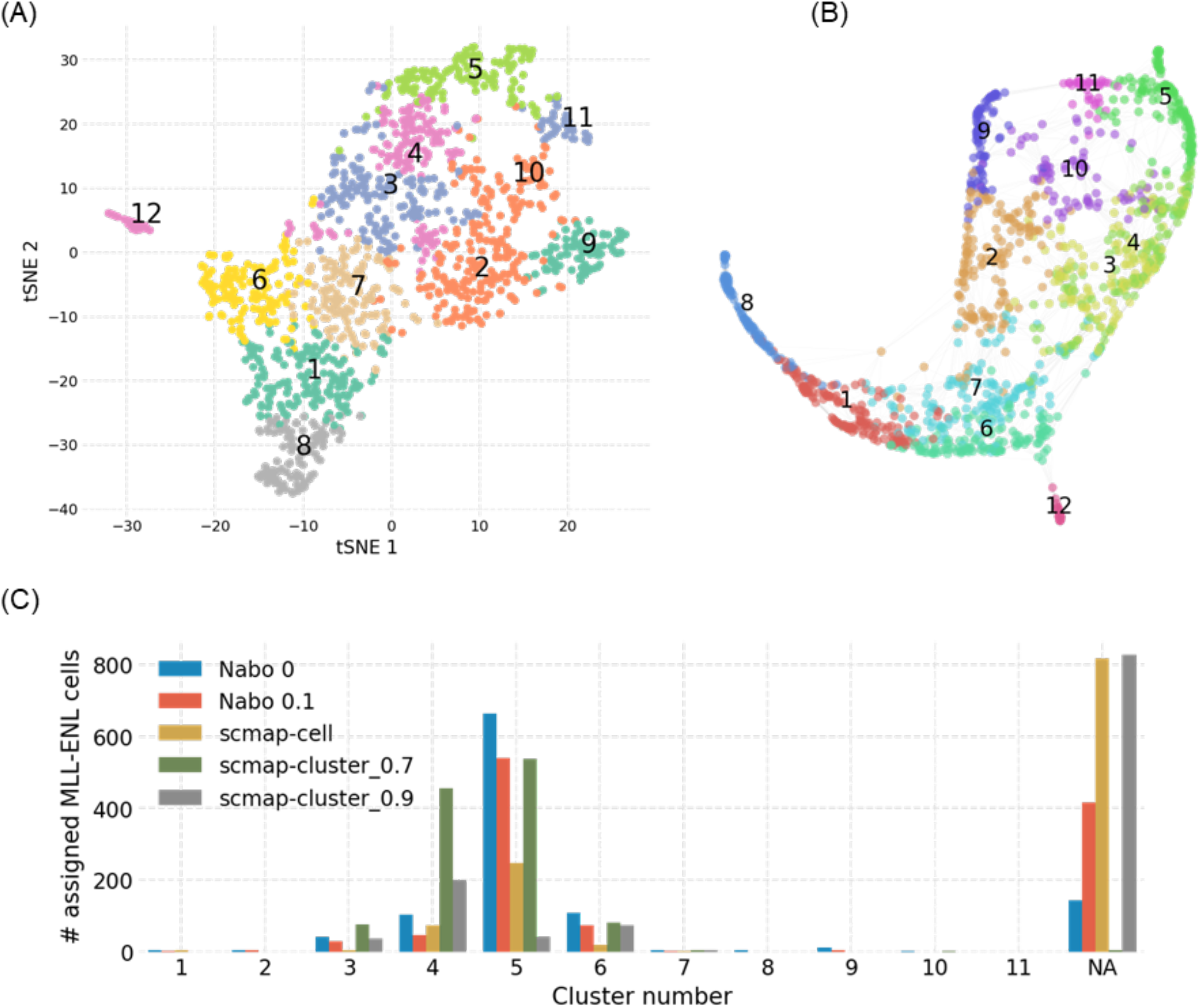
(**A**) A t-SNE plot of normal GMLPs obtained using Seurat. The cells have been colored based on cluster association, which is also indicated by numerals. (**B**) The SNN graph (from Figure 4A) of normal GMLPs, using the same cluster identities as shown in A. (**C**) Barplots showing the number of MLL-ENL cells that were assigned to each cluster obtained from Seurat. The minimum weight for assignment using Nabo was set at either 0 or 0.1 (default). In case of scmap-cluster the weight fraction was set either at 0.7 (default) or 0.9.

**Supplementary figure 4:**
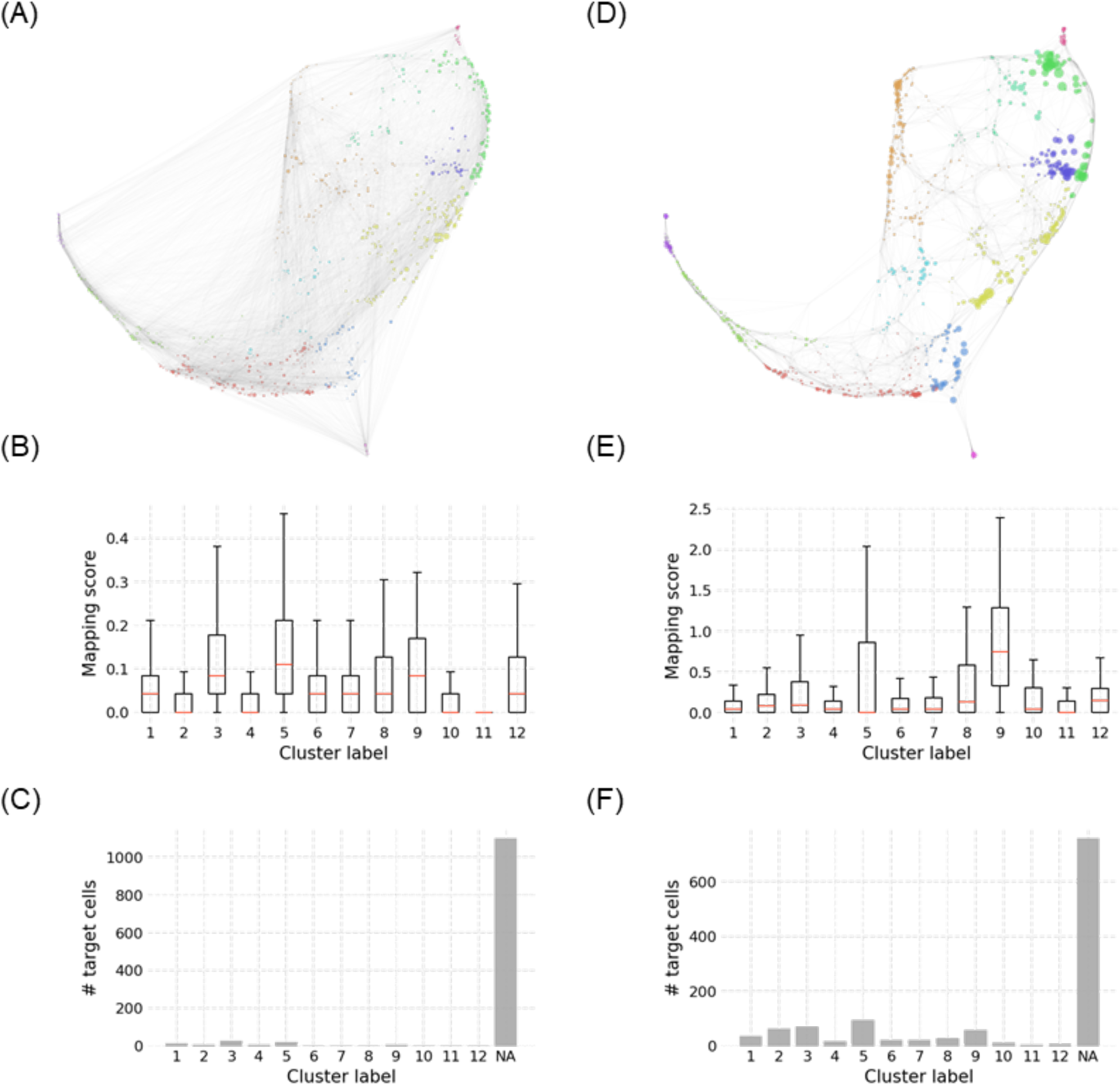
(**A**) SNN graph of normal GMLPs constructed using low variance genes sampled from the same expression range as the highly variable genes (HVGs). The cell positions have been set to be the same as in the SNN graph obtained using the HVGs (**figure 4A**). The cell size has been scaled to indicate the mapping score obtained when MLL-ENL induced GMLPs were projected on this graph. (**B**) The cluster-wise distribution of mapping scores shown in A. The same cluster identities were used in the HVG graph. (**C**) The number of MLL-ENL cells assigned to each of the cluster. (**D**) SNN graph of WT GMLPs following blocked mapping. (**E**) The cluster-wise distribution of mapping scores shown in D. The same cluster identities were used in the HVG graph. (**F**) The number of MLL-ENL cells assigned to each of the cluster after the projection shown in D.

**Supplementary figure 5:**
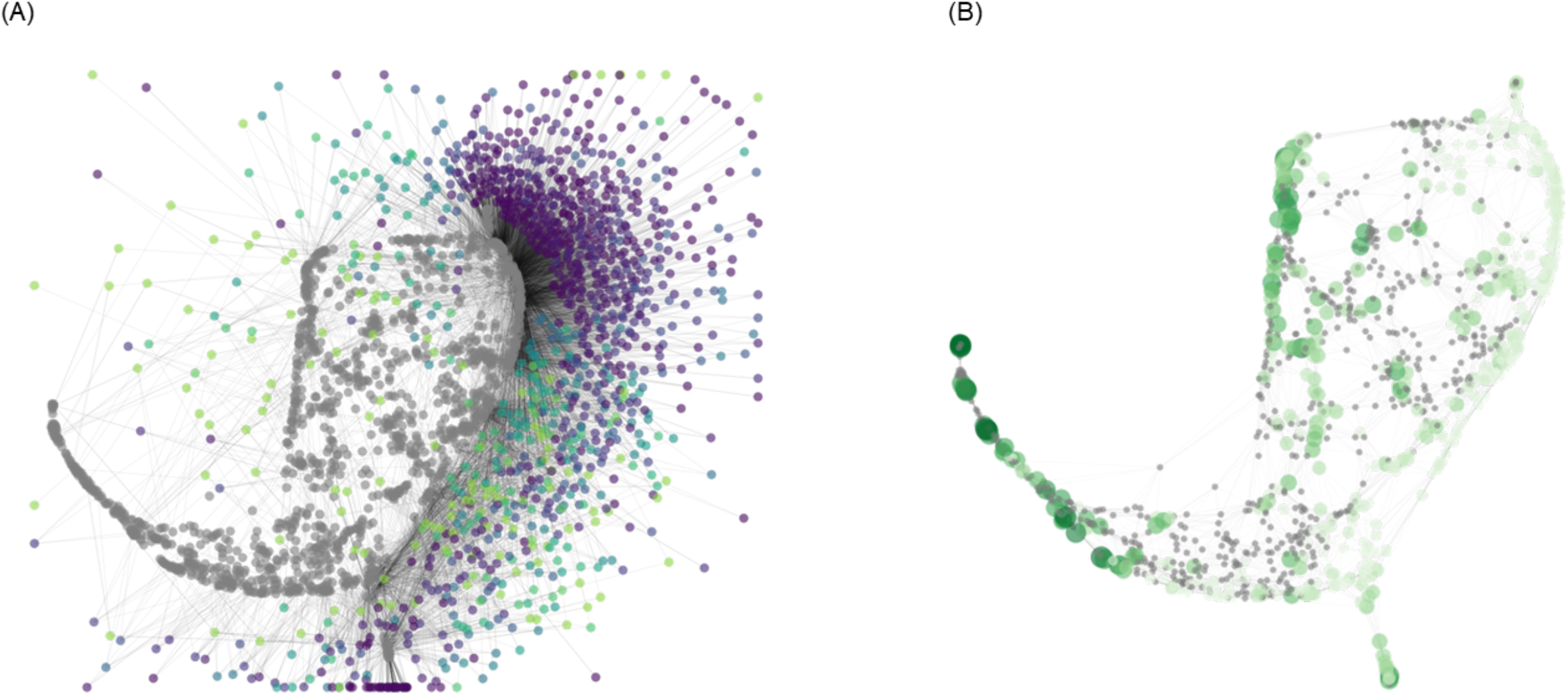
(**A**) Force directed layout of SNN graph of normal GMLPs along with the force directed layout of MLL-ENL cells. The reference cells are in gray and the MLL-ENL cells are colored based on their mapping specificity. The mapping specificity is indicated in dark purple (highest) and light yellow (least specificity). (**B**) The average mapping specificity of each reference cell based on the specificity of each MLL-ENL cell that mapped onto those reference cells. Smaller and lighter colored cells have higher mapping specificies and those in large size and dark green color have lower specificity. The cells shown in grey were not mapped to.

**Supplementary figure 6:**
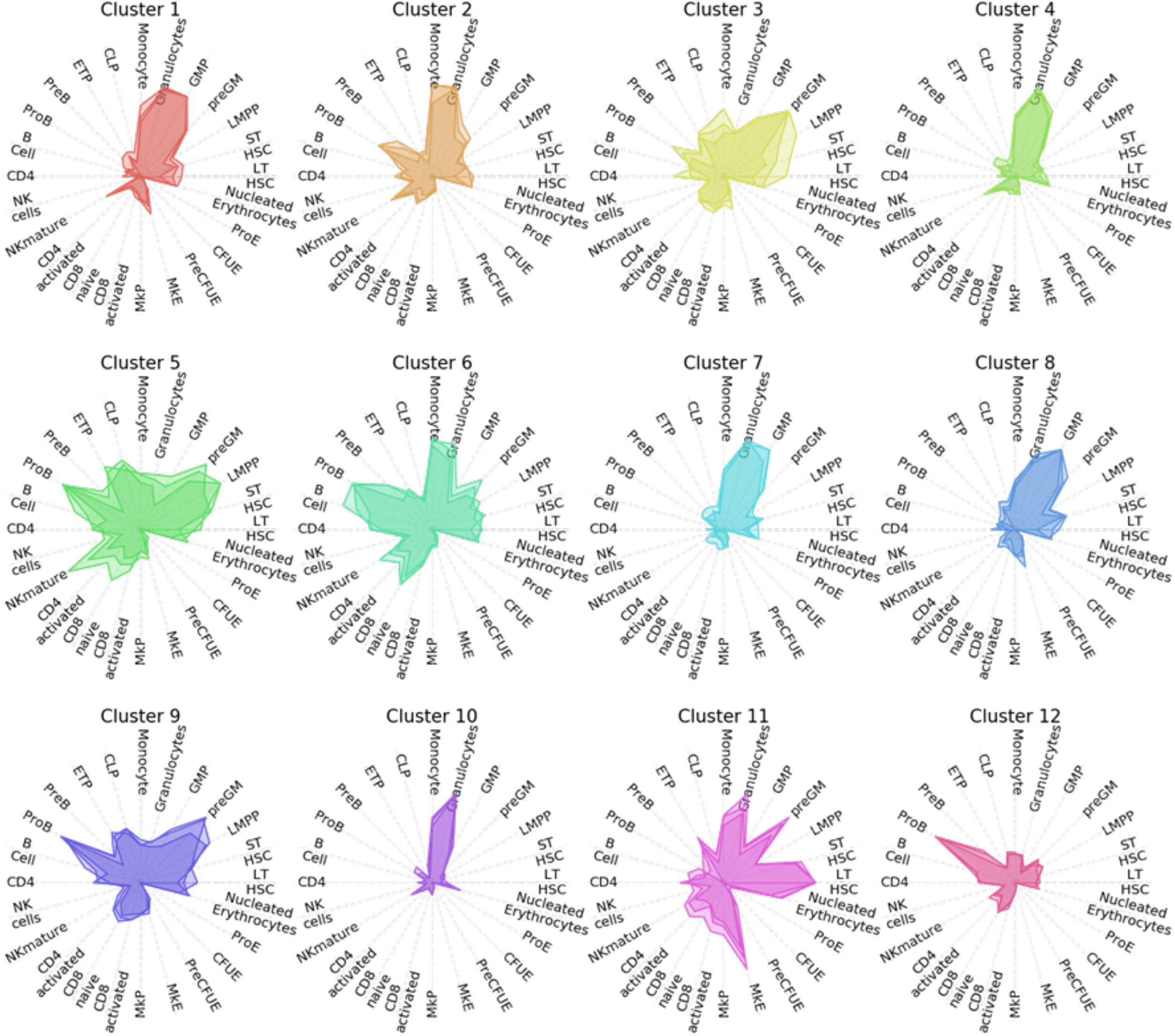
(**A**) Radar plots showing the predicted cell type for each of the normal GMLP cluster. The three area polygons in the radar plot show the mean and +/− 1 SD values.

**Supplementary figure 7:**
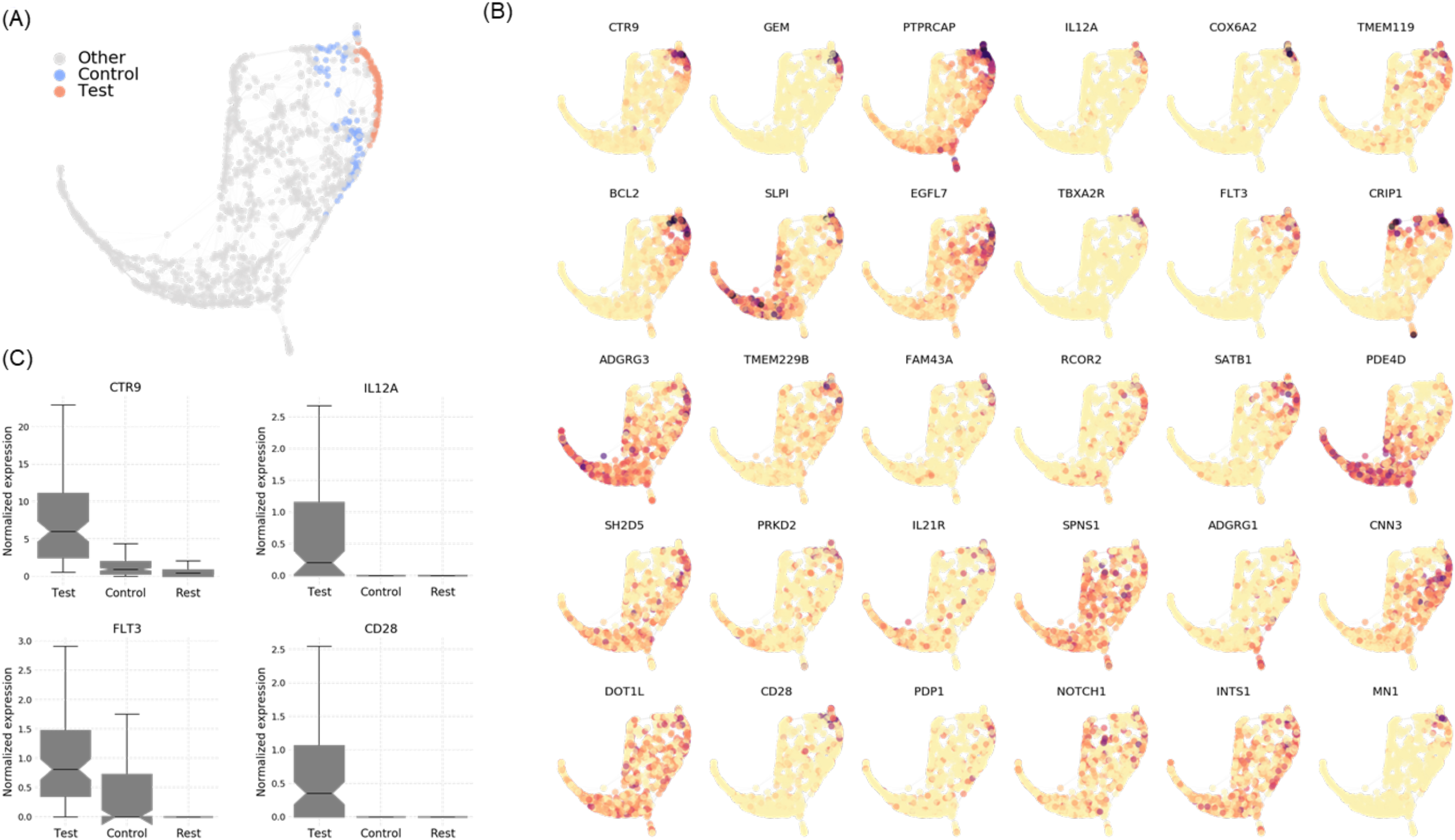
(**A**) Force directed SNN graph of normal GMLPs. The red dots represent cells with mapping scores higher than 1 and are from cluster 5. The blue dots represent cells which are at a path distance of 2 from red cells. The red cells were compared with blue cells to identify differentially expressed genes associated with mapped cells. (**B**) The expression of the top 30 most differentially expressed genes. The cells marked in dark red have higher expression and the ones marked in yellow have the less expression. (**C**) Notched boxplots showing the distribution of selected genes in the three groups.

**Supplementary figure 8:**
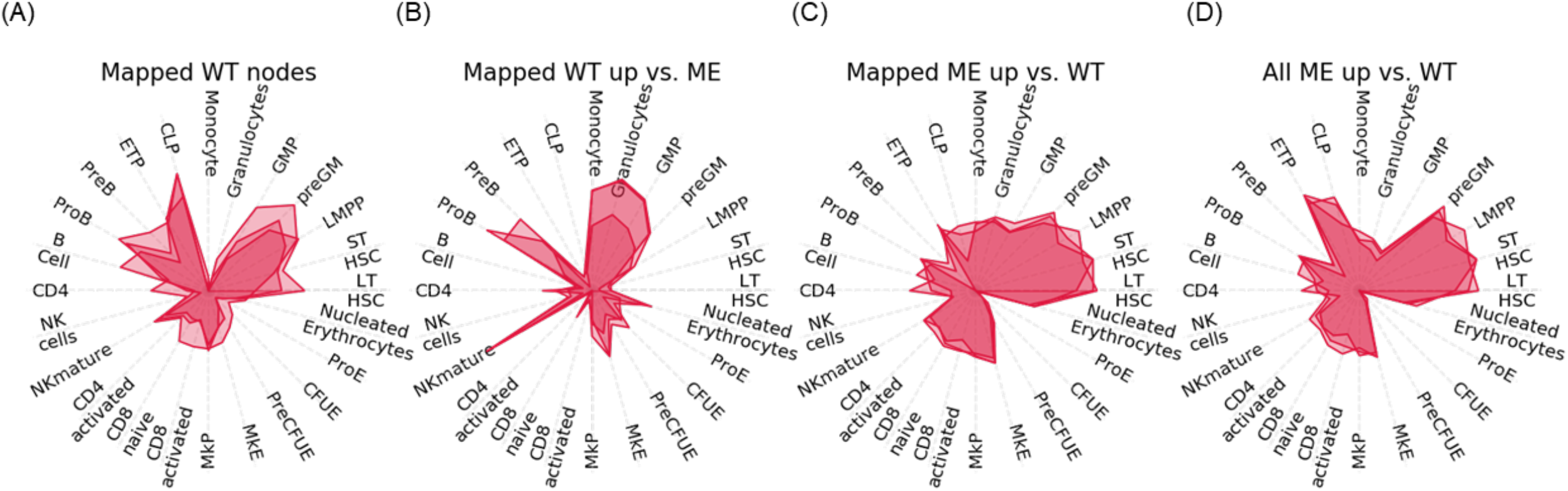
Radar plots showing the gene expression-based lineage affiliation of cells. Genes used to generate each plot were those differentially expressed (upregulated) obtained by comparing: (**B**) Normal GMLPs from cluster 5 that received mapping scores over 1 compared to other cells at path distance of 2. (**B**) WT GMLPs from cluster 5 that received mapping scores over 1 compared to MLL-ENL cells that mapped to these cells. (**C**) The opposite approach of B. (**D**) All MLL-ENL cells compared to all normal GMLPs.

**Supplementary figure 9:**
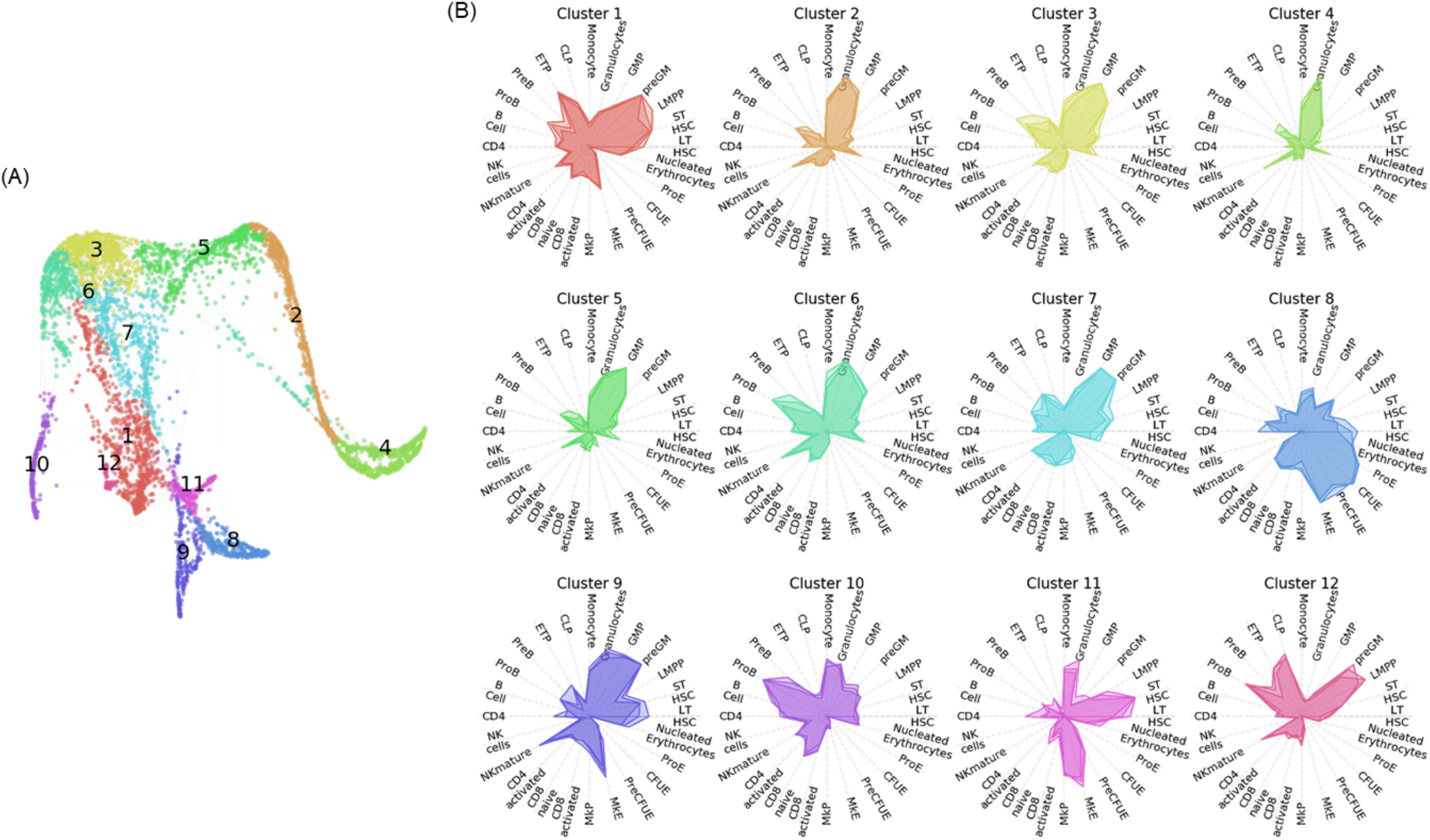
(**A**) SNN graph of c-kit+ cells, where cells have been colored based on the cluster identity. (**B**) Radar plots showing the lineage bias of each cluster, inferred from gene expression signature of each cluster.

## ACKNOWLEDGMENTS

We thank Mikael Sommarin, Johan Rodhe and Oscar Legetth at Lund Stem Cell Center for their help with code and documentation review. This work was supported by grants from the Swedish Cancer Society, The Ragnar Söderberg Foundation, the Knut and Alice Wallenberg Foundation, the Swedish Research Council, the Swedish Society for Medical Research, and the Swedish Childhood Cancer Foundation.

## AUTHOR CONTRIBUTIONS

G.K, D.B, and P.D conceived and designed the study; D.B, A.U, M.E, and E.E designed and performed mouse experiments and sequencing; P.D and S.S designed and performed the bioinformatics- and computational analyses; P.D and R.O packaged Nabo codebase and prepared documentation; G.K, D.B, and P.D analyzed and interpreted data; G.K and P.D prepared the figures and wrote the manuscript with input from all authors.

